# AP-3 and the V-ATPase Modulate CTP Synthase Assembly Through Spatial Association at the Yeast Vacuole

**DOI:** 10.64898/2026.02.13.705788

**Authors:** Michaela McCright, Mitchell Leih, Ayla Nack, Cortney G. Angers, Alexey J. Merz, Greg Odorizzi

**Author notes:** Address correspondence to: Greg Odorizzi.

## Abstract

The compartmentalization of metabolic enzymes into membraneless filaments termed cytoophidia represents a conserved regulatory mechanism, exemplified by cytidine triphosphate synthase (CTP synthase). CTP synthase assembles into pH-sensitive cytoophidia in the cytosol. In *Saccharomyces cerevisiae*, nutritional deprivation both triggers CTP synthase cytoophidia assembly and disassembles the vacuolar H□-ATPase (V-ATPase) that acidifies vacuoles (lysosomes), yet whether these processes are functionally linked remains unknown. We demonstrate spatial proximity between the yeast CTP synthase homolog Ura7, the V-ATPase, and the AP-3 adaptor complex that mediates vesicular transport to vacuoles. We show Ura7 localizes to vacuoles under both nutrient-rich and starvation conditions. Genetic disruption of AP-3 function altered Ura7 assembly dynamics under starvation, reducing total structures yet dramatically enhancing Ura7 cytoophidia elongation (∼5-fold), revealing a dual regulatory role for AP-3 that both promotes Ura7 assembly and restrains elongation. Moreover, combining nutritional and pharmacological V-ATPase inhibition triggered massive Ura7 cytoophidia formation. These findings reveal a previously unrecognized spatial coupling between metabolic enzyme compartmentalization, vacuolar trafficking, and the pH regulation machinery, suggesting a new organizational principle whereby CTP synthase assembly dynamics respond to vacuolar function.

## INTRODUCTION

The AP-3 (Adaptor Protein complex-3) pathway mediates vesicular transport from late Golgi/endosomal compartments to lysosomes and lysosome-related organelles (Cowles et al., 1997; Theos et al., 2005; Robinson, 2015). Unlike AP-1 and AP-2 complexes, AP-3 remains on coated vesicles during transit, releasing only upon arrival at their target organelle where they facilitate tethering through the HOPS complex (Angers and Merz, 2009; Cabrera et al., 2010; Schwartz et al., 2017; Schoppe et al., 2020). This prolonged coat association may enable spatial organization of cytosolic factors to sites of membrane trafficking and fusion. AP-3 proximity-labeling revealed numerous proteins beyond canonical AP-3 cargoes that reside in close spatial proximity to this trafficking machinery (Schoppe et al., 2020), raising questions about whether AP-3 coordinates additional cellular processes beyond vesicle transport. Among these proximity-detected proteins are metabolic enzymes, suggesting potential links between vesicular trafficking and metabolic regulation.

Cytidine triphosphate synthase (CTP synthase) is one of the metabolic enzymes identified in the AP-3 proximity dataset (Schoppe et al., 2020). CTP synthase exemplifies an emerging regulatory mechanism through self-assembly into membraneless filaments termed cytoophidia (Noree et al., 2010; Liu, 2016). In *Saccharomyces cerevisiae*, the CTP synthase homolog Ura7 forms cytoophidia in response to stressors including glucose starvation, which sequesters the enzyme in an inactive state (Barry et al., 2014; Noree et al., 2014). Ura7 cytoophidium formation is exquisitely sensitive to intracellular pH, with acidification triggering rapid assembly (Hansen et al., 2021). This pH sensitivity suggests potential coordination with the vacuolar H□-ATPase (V-ATPase), which regulates both organellar and cytosolic pH through reversible assembly of cytoplasmic (V1) and membrane (V0) sectors (Kane, 2006; Martinez-Munoz and Kane, 2008; Forgac, 2007). Notably, glucose starvation—the same condition that induces Ura7 cytoophidia—triggers rapid V1-V0 disassembly, silencing proton transport and causing cytosolic acidification (Kane, 1995; Parra et al., 2000).

Despite extensive characterization of AP-3 trafficking, CTP synthase assembly dynamics, and V-ATPase regulation, how these three systems—membrane trafficking, metabolic enzyme compartmentalization, and pH homeostasis—might be functionally integrated remains unknown. The convergent observation that glucose starvation both disassembles V-ATPase and induces Ura7 cytoophidia (Kane, 1995; Noree et al., 2010) hints at coordination, but direct connections have not been demonstrated. Intriguingly, multiple V-ATPase subunits appear in an AP-3 proximity-labeling dataset (Schoppe et al., 2020) as well as in our recent AP-3 affinity chromatography analysis (Leih et al., submitted), and V-ATPase mutants exhibit AP-3 trafficking defects (Anand et al., 2009), suggesting that the trafficking machinery and pH regulatory apparatus may associate. Whether this proximity extends to metabolic enzymes like CTP synthase remains unexplored.

By systematically interrogating published AP-3 interactomes in yeast using bimolecular fluorescence complementation (BiFC), we discover spatial associations between the CTP synthase Ura7 and both AP-3 and the V-ATPase. Genetic ablation of AP-3 sensitizes Ura7 to stress-induced filamentation, and V-ATPase inhibition using both nutritional and pharmacological methods triggers massive Ura7 assembly, resulting in extensive cytoophidia formation. These findings reveal previously unrecognized connections between the vacuolar trafficking machinery, pH regulatory apparatus, and metabolic enzyme compartmentalization.

## RESULTS AND DISCUSSION

### Spatial association between AP-3 and CTP synthases

To identify candidate AP-3-associated proteins in yeast, we leveraged proteomics datasets from AP-3 proximity-labeling (Schoppe et al., 2020) and our recent AP-3 affinity chromatography study (Leih et al., submitted). Ninety-eight proteins were identified by both methods (Supplemental Table S1), providing a focused validation list that we screened for AP-3 association *in vivo* by BiFC. We recently demonstrated that BiFC can characterize known AP-3 associations and reveal novel ones (Leih et al., submitted). We tested candidates using an *S. cerevisiae* VN fusion library (Sung et al., 2013), identifying 6 positives showing BiFC signal when coexpressed with a VC fusion to the Apl6 large subunit of AP-3 (Supplemental Figure S1). Among these, we focused on Ura7 (CTP synthase), motivated by its glucose starvation-induced cytoophidium formation (Noree et al., 2010), a condition that also inhibits the V-ATPase (Kane, 1995; Parra and Kane, 1998; ).

Given that AP-3 vesicles transport cargo proteins to the vacuole, we hypothesized Ura7 proximity with AP-3 might link nucleotide metabolism to vacuolar trafficking under nutrient stress. To assess whether this spatial association involves the assembled AP-3 complex, we also tested for BiFC between Ura7 and the Apl5 large subunit of AP-3. As shown in Figure 1A, BiFC puncta appeared when Ura7-VN was paired with Apl6-VC or Apl5-VC but not with the negative-control His2-VC fusion. Both Apl6 and Apl5 pairings with Ura7 showed significantly more BiFC-positive cells than the His2 control (Figure 1B), and quantification of total BiFC fluorescence per cell showed similar trends for both AP-3 subunits (Figure 1C). Apl6-VC and Apl5-VC also produced BiFC puncta with Ura8-VN (Supplemental Figure S2). Ura7 and Ura8 are 78% identical and are functionally overlapping enzymes with differential regulation—Ura7 being the major isoform (Ozier-Kalogeropoulos et al., 1994).

**Figure 1.**
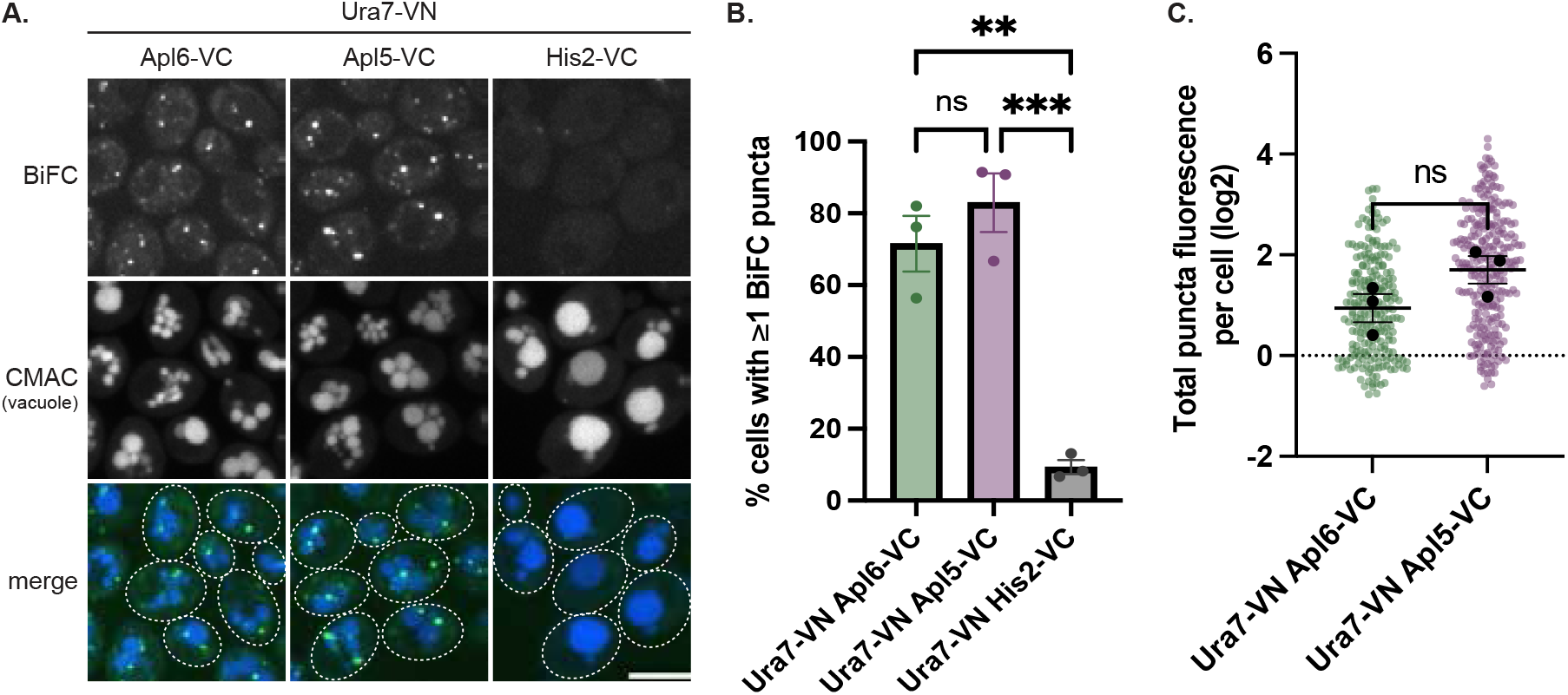
BiFC validates spatial proximity between AP-3 and Ura7. **(A)** Representative confocal images of cells expressing the indicated VN/VC fusions. Scale bar, 5 μm. **(B)** Quantification of BiFC-positive cells (percentage of cells with ≥1 BiFC punctum); circles = biological replicate means (n=3, >100 cells/replicate), error bars = SEM. Statistical comparisons by one-way ANOVA with Tukey’s post-hoc test; **p=0.0013; ***p=0.0005; ns, not significant (p=0.481). **(C)** Quantification of total BiFC fluorescence per cell (log-transformed integrated density) for Apl6-VC/Ura7-VN and Apl5-VC/Ura7-VN. Black circles = biological replicate means (n=3, >100 cells/replicate), colored circles = individual cell measurements, error bars = SEM. Comparison by unpaired t-test; ns, not significant (p=0.122). .

The identification of Ura7 and Ura8 CTP synthases in AP-3 proximity datasets places these redundant metabolic enzymes near the Golgi-to-vacuole trafficking pathway, suggesting spatial metabolic regulation. AP-3 vesicle formation at the Golgi and tethering at the vacuole (Rehling et al., 1999; Angers and Merz, 2009) may position CTP synthases near organelle surfaces, potentially linking nucleotide synthesis with trafficking events. Because both AP-3 large subunits show comparable BiFC with both CTP synthase isoforms, their spatial association involves the assembled AP-3 complex. This dual recognition mirrors the redundancy in cytoophidium formation, where Ura7 and Ura8 colocalize (Noree et al., 2010). Neither Ura7 nor Ura8 contain canonical AP-3 sorting signals, and the spatial proximity we observe is distinct from cargo-coat interactions driving selective transport. Rather, CTP synthases may localize to sites where AP-3 vesicles interact with the vacuole.

### BiFC reveals spatial associations linking AP-3, Ura7, and the V-ATPase

The BiFC signal between AP-3 and CTP synthases suggested a connection between AP-3 and cellular metabolism. As noted above, Ura7 and Ura8 form cytoophidia in response to glucose starvation (Noree et al., 2010), a metabolic stress that also inhibits the V-ATPase (Kane, 1995, Parra and Kane, 1998). This prompted us to consider if AP-3 might also be spatially associated with V-ATPase components. AP-3 proximity labeling had identified six V-ATPase subunits (Schoppe et al., 2020), among which, Vma13 was detected in our AP-3 affinity chromatography study (Leih et al., submitted; Supplemental Table S1). We observed no BiFC signal between Vma13 and Apl6/Apl5, though this negative result might reflect the essential nature of the C-terminal domain of Vma13 where VN was fused (Liu et al., 2005). We therefore tested Vma10, which is a V-ATPase subunit positioned near Vma13 (Forgac, 2007) and was among the subunits identified by AP-3 proximity labeling (Schoppe et al., 2020). Cells expressing Vma10-VN with either Apl6-VC or Apl5-VC showed BiFC puncta adjacent to vacuoles, while His2-VC/Vma10-VN controls showed minimal signal (Figure 2A). Both Apl6 and Apl5 pairings with Vma10 showed significantly more BiFC-positive cells than His2 controls (Figure 2B), and total BiFC fluorescence per cell showed similar trends for both AP-3 subunits (Figure 2C).

**Figure 2.**
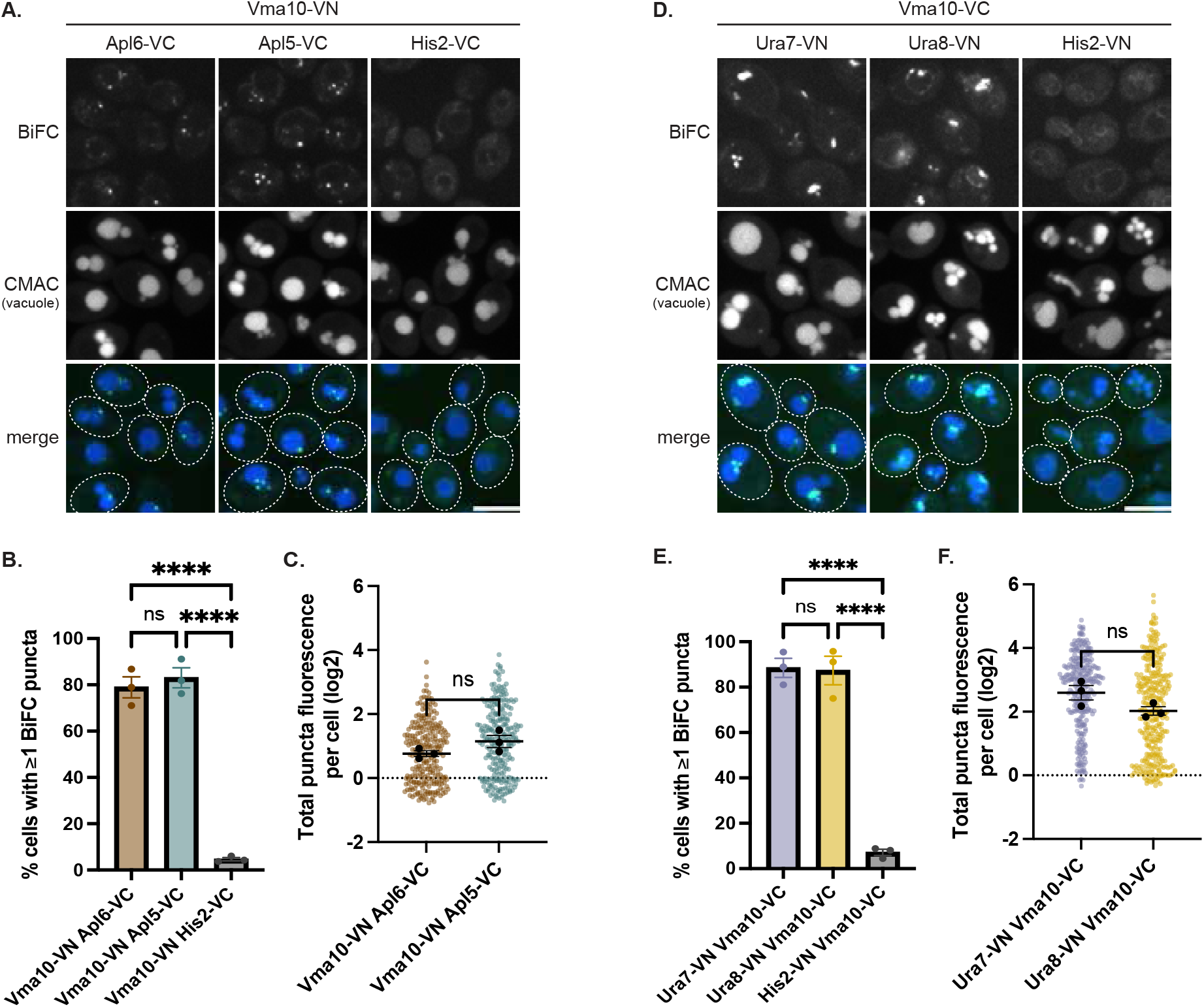
AP-3 and Ura7 associate with the V-ATPase subunit Vma10. Representative confocal images of cells expressing the indicated VN/VC fusions **(A**,**D)**. Scale bar, 5 μm. Quantification of **(B)** BiFC-positive cells; ****p<0.0001; ns, not significant (p=0.719) and **(C)** total BiFC fluorescence; ns, not significant (p=0.148). Graphs and statistics as in Figure 1B,C. Quantification of **(E)** BiFC-positive cells; ****p<0.0001; ns, not significant (p=0.980) and **(F)** total BiFC fluorescence; ns, not significant (p=0.095). Graphs and statistics as in Figure 1B,C.

Given AP-3 proximity to both Ura7 and Vma10, we tested by BiFC whether Ura7 and Vma10 are also in close proximity. Indeed, cells expressing Vma10-VC with either Ura7-VN or Ura8-VN showed BiFC puncta, while the Vma10-VC/His2-VN control showed minimal signal (Figure 2D). Both Ura7 and Ura8 pairings with Vma10 showed significantly more BiFC-positive cells than His2 controls (Figure 2E), and total fluorescence measurements showed similar trends (Figure 2F). Consistent with this association, independent proteomic analysis identified Ura7 in immunoprecipitates of the V-ATPase from cell extracts (Khan et al., 2021). These spatial associations suggest AP-3, CTP synthases, and the V-ATPase function within a spatially organized network coordinating nucleotide metabolism and vacuolar acidification.

The V-ATPase itself does not traffic to vacuoles via the AP-3 pathway (Piper et al., 1997; Bowers et al., 2005). Thus, the presence of V-ATPase subunits in the AP-3 proximity cloud supports AP-3 coat retention until vesicles reach the vacuole (Angers and Merz, 2009; Schoppe et al., 2020). Vma10 is a peripheral stalk component connecting the V1 and V0 sectors of the V-ATPase (Charsky et al., 2000; Kane, 2006; Ohira et al., 2006). CTP synthase is enzymatically inactive when assembled into cytoophidia (Barry et al., 2014), and the formation of these filaments is triggered by drops in intracellular pH (Hansen et al., 2021). Because the V-ATPase is the primary regulator of cytosolic and organellar pH (Martinez-Munoz and Kane, 2008), spatial proximity between Vma10 and CTP synthases could provide a mechanism for CTP synthase assembly to respond to the V-ATPase structural state.

### Ura7 localizes to vacuoles under nutrient-rich and starvation conditions

We next examined whether the spatial associations between Ura7, AP-3, and the V-ATPase reflect functional localization at the vacuole. While cytoophidium formation increases upon glucose starvation, basal CTP synthase assembly into punctate structures (foci) has been observed in unstressed cells (Noree et al., 2010). If Ura7 coordinates with AP-3 and the V-ATPase, Ura7 structures should localize to the vacuolar membrane where these complexes reside. We examined cells coexpressing Ura7-GFP and Vph1-mCherry (V-ATPase vacuolar marker) in rich medium versus nutrient starvation (citrate-phosphate buffer, overnight). Ura7-GFP events (defined as Ura7-GFP assembled either as punctate structures or as filamentous cytoophidia) were infrequent in rich medium, consistent with low basal assembly in unstressed cells, but more than one-third of these overlapped Vph1-mCherry at the vacuolar membrane (Figure 3A,D). After starvation, the number of events increased more than two-fold (Figure 3B,C), but a similar proportion of the total events overlapped Vph1-mCherry at vacuoles (Figure 3D). Consistent with this localization, an unbiased proteomic analysis of purified vacuoles identified Ura7 as a vacuolar protein (Eising et al., 2022). Collectively, these observations indicate that Ura7 localizes to the vacuolar membrane regardless of metabolic state, with the extent of assembly being modulated by nutrient availability.

**Figure 3.**
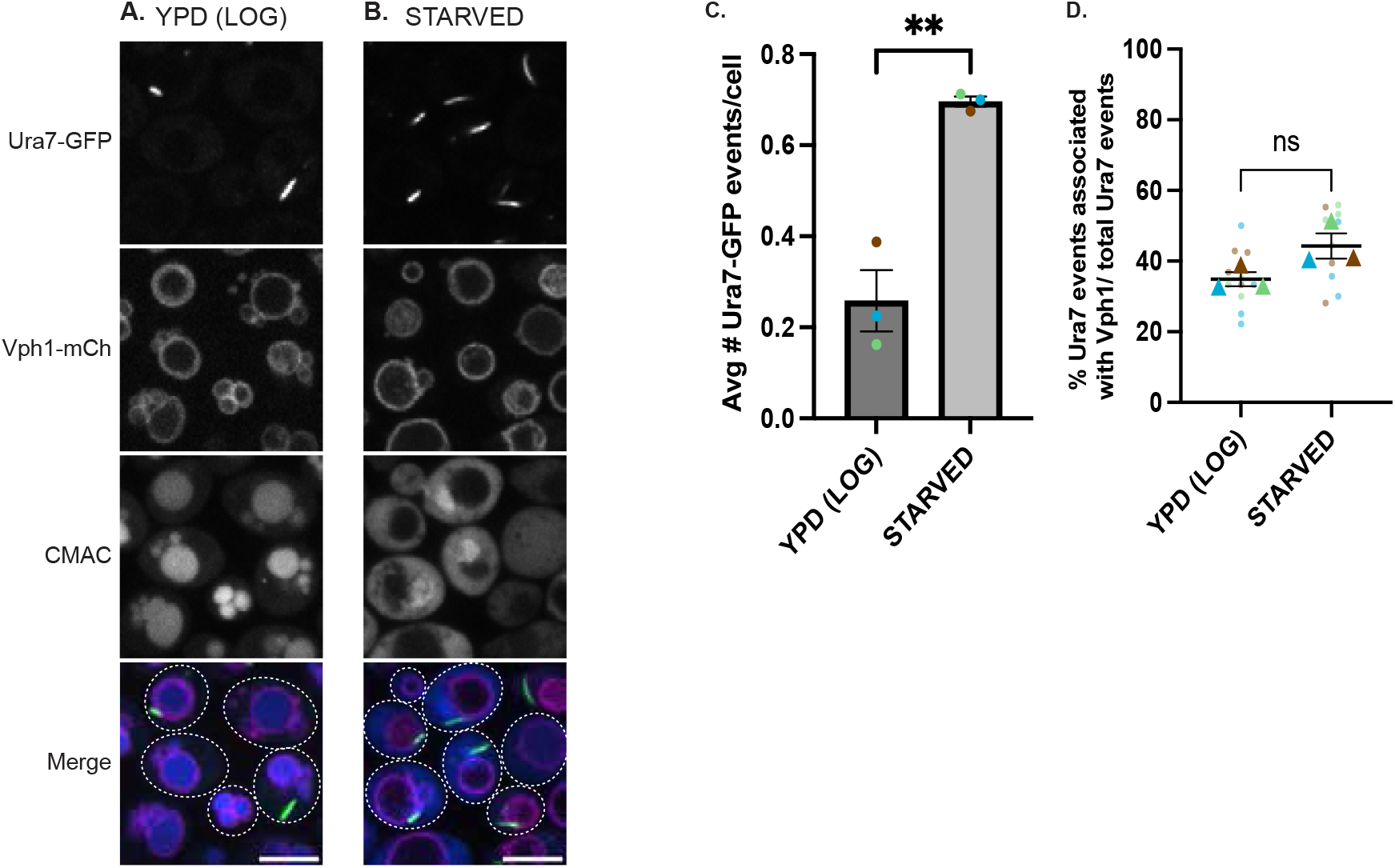
Ura7 cytoophidia localize to the vacuole under both nutrient-rich and starvation conditions. Representative confocal images of cells expressing Ura7-GFP and Vph1-mCherry in rich medium **(A)** or starvation medium **(B)**. Scale bar, 5 μm. CMAC-blue marks the vacuolar lumen in rich medium but marks the cytosol in starvation medium. **(C)** Quantification of average Ura7-GFP events (puncta or cytoophidia) per cell in rich vs starvation conditions. **p=0.003 by unpaired t-test. Brown, blue, and green represent the average of each replicate. **(D)** Quantification of Ura7-GFP events colocalizing with Vph1-mCherry (% of total events). Object-based colocalization analysis described in Materials and Methods. ns, not significant (p=0.084) by unpaired t-test. Brown, blue, and green triangles represent the average of each replicate.

### AP-3 loss promotes Ura7 cytoophidia elongation under starvation

Having established that Ura7 localizes to vacuoles, we next examined whether AP-3 function affects Ura7 assembly dynamics using Ura7-mCherry, which exhibits low basal assembly in unstressed cells, facilitating detection of starvation-induced changes. We examined Ura7-mCherry in wild-type, *apl5*_Δ_, and *apl6*_Δ_ cells cultured at log phase versus starvation. Log-phase cells showed diffuse cytosolic/nuclear Ura7-mCherry across all three strains with minimal punctate or filamentous cytoophidia structures (Figure 4A-C). After starvation, wild-type cells showed a mixture of puncta and cytoophidia, while AP-3 mutants dramatically shifted toward elongated cytoophidia, increasing the filament-to-puncta ratio ∼5-fold (Figure 4G). Ratios in *apl5*_Δ_ and *apl6*_Δ_ were indistinguishable from one another, consistent with complete loss of AP-3 complex function. Interestingly, AP-3 mutants showed significantly fewer total Ura7-mCherry events per cell compared to wild-type (Figure 4H), indicating that AP-3 function promotes Ura7 assembly. Together, these results reveal a dual role for AP-3 in modulating Ura7 assembly dynamics under starvation: AP-3 promotes the formation of Ura7 structures while simultaneously restraining their elongation into extended cytoophidia. The dramatic shift in filament-to-puncta ratio despite reduced total events suggests that, once nucleated in the absence of AP-3, Ura7 structures more readily mature into elongated cytoophidia.

**Figure 4.**
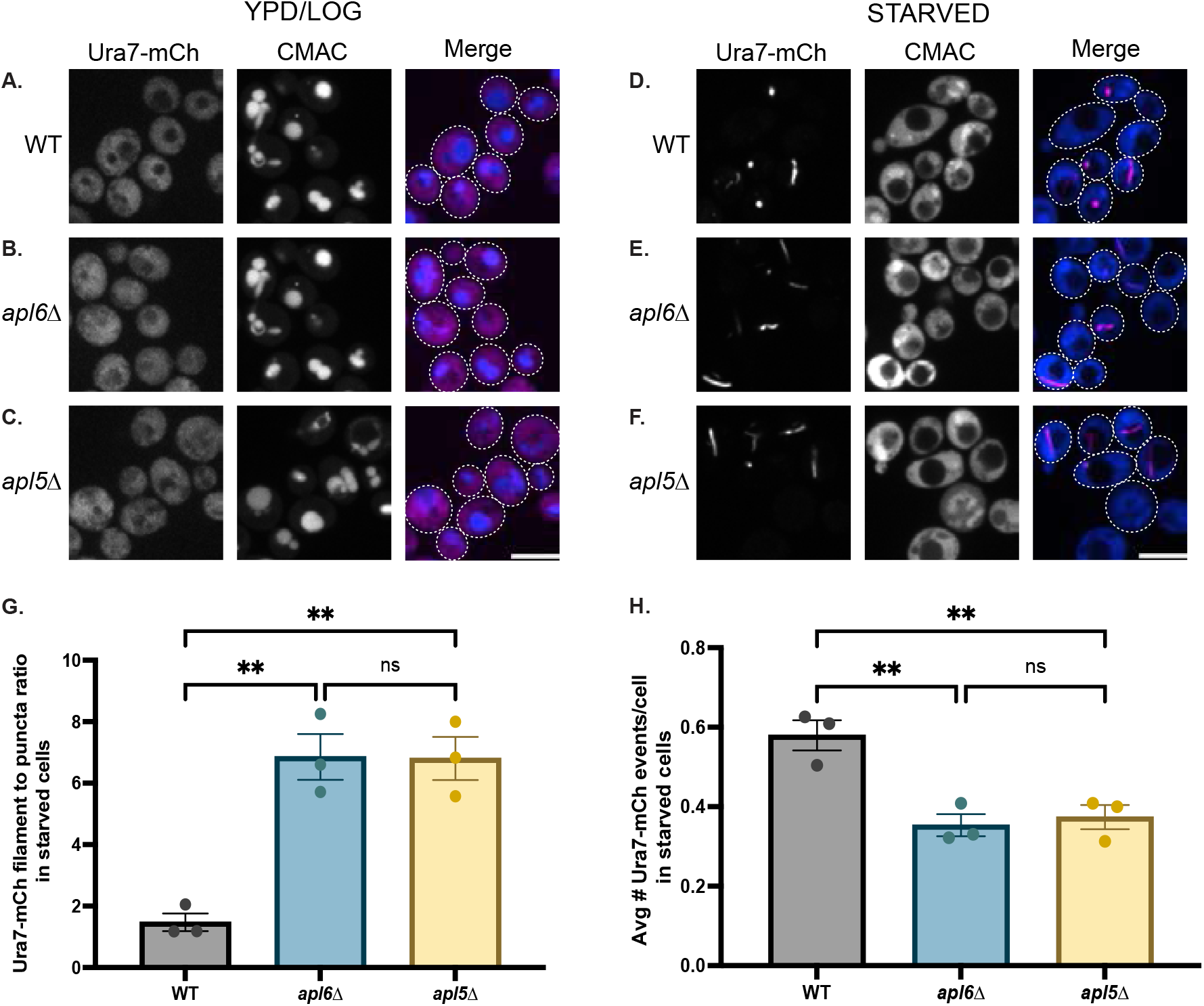
Loss of AP-3 function promotes Ura7 hyper-assembly under starvation conditions. Representative confocal images of wild-type **(A**,**D)**, *apl6Δ* **(B**,**E)**, and *apl5Δ* **(C**,**F)** cells expressing Ura7-mCherry in rich medium (A-C) or after starvation (D-F). Scale bar, 5 μm. CMAC-blue staining as in Figure 3. **(G)** Quantification of Ura7-mCherry filament-to-puncta ratio in starved cells (filaments = aspect ratio ≥1.5, puncta = aspect ratio <1.5). **p=0.002 vs WT; ns (p=0.998) between *apl6Δ* and *apl5Δ* by one-way ANOVA with Tukey’s post-hoc test. **(H)** Quantification of total Ura7-mCherry events per cell in starved cells. Statistics as in (G). *apl6Δ* vs WT: **p=0.006; *apl5Δ* vs WT: **p=0.0098; *apl6Δ* vs *apl5Δ*: ns, not significant (p=0.899).

The opposing effects of AP-3 loss—reduced event frequency yet enhanced cytoophidia elongation—suggest AP-3 influences multiple aspects of Ura7 regulation. An independent proteomic analysis of purified vacuoles found Ura7 is depleted from vacuoles in AP-3 mutants (Eising et al., 2022), consistent with our observation that AP-3 mutants show fewer total Ura7 events. These results suggest that in wild-type cells, a pool of Ura7 associates with vacuoles in an AP-3-dependent manner. The loss of this vacuolar-associated pool in AP-3 mutants, combined with the cellular stress of impaired vacuolar trafficking (Darsow et al., 1998; Sun et al., 2004; LaGrassa and Ungermann, 2005), could lower the threshold for filament assembly, but the specific mechanism by which AP-3 loss enhances cytoophidia elongation remains to be determined. Since Ura7 cytoophidia formation is triggered by cytosolic acidification (Hansen et al., 2021), pH dysregulation could contribute, though we have not measured pH in AP-3 mutants. Alternative mechanisms—including altered metabolic signaling or general stress responses—warrant investigation.

### V-ATPase inhibition induces Ura7 assembly and cytoophidia formation

We next asked whether V-ATPase inhibition affects Ura7 assembly dynamics. The V-ATPase can be inhibited by glucose withdrawal, which triggers V1-V0 disassembly (Kane, 1995), or by concanamycin A (ConcA), which blocks proton translocation (Dröse and Altendorf, 1997). We subjected wild-type cells expressing Ura7-mCherry to four conditions: mock treatment, glucose withdrawal, ConcA treatment, or combined treatment (see Materials and Methods for details). While mock-treated cells showed diffuse cytosolic Ura7-mCherry (Figure 5A), glucose withdrawal (Figure 5B) or ConcA treatment (Figure 5C) induced Ura7 to transition into aggregates with some filaments. Strikingly, the combined treatment produced a dramatically enhanced phenotype, with marked increases in both the number and size of puncta plus extensive elongated cytoophidia formation (Figure 5D-H). The number of filamentous structures (aspect ratio >1.5 and area >0.5 μm^2^) was substantially elevated compared to either single treatment alone (Figure 5H), demonstrating that dual V-ATPase inhibition potently induces Ura7 cytoophidia elongation.

**Figure 5.**
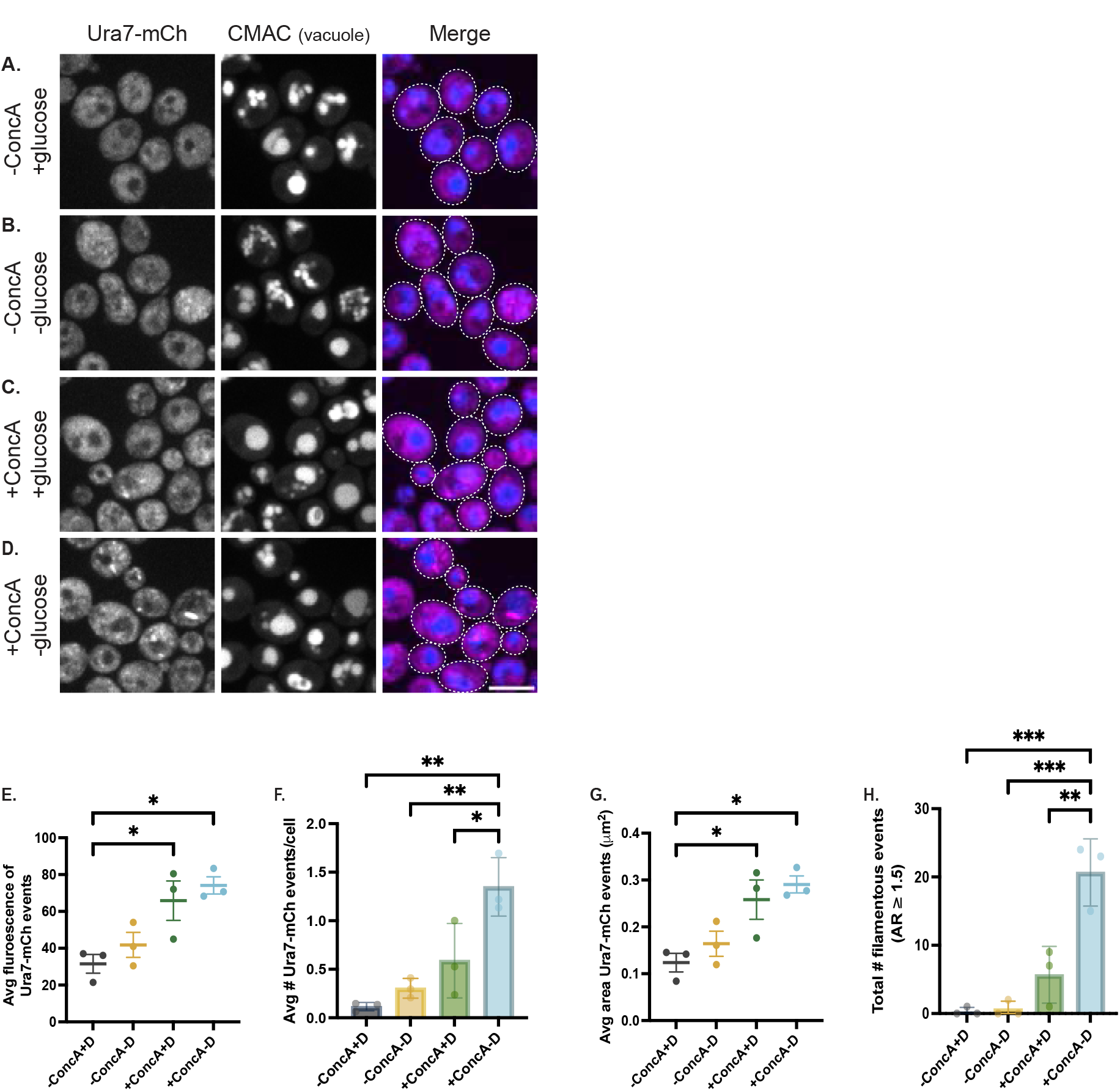
Dual inhibition of the V-ATPase potently induces Ura7 assembly and filament formation. Representative confocal images of wild-type cells expressing Ura7-mCherry subjected to: **(A)** mock treatment, **(B)** glucose withdrawal, **(C)** ConcA treatment, or **(D)** dual treatment. Scale bar, 5 μm. **(E)** Quantification of average fluorescence intensity (integrated density) of Ura7-mCherry structures per cell; circles = biological replicate means (n=3, >100 cells/replicate), error bars = SEM. Statistics by one-way ANOVA with Tukey’s post-hoc test. -ConcA+D vs +ConcA+D: *p=0.04; -ConcA+D vs +ConcA-D: *p=0.013; all other comparisons not significant. **(F)** Quantification of Ura7-mCherry events per cell. Statistics as in (E). -ConcA+D vs +ConcA-D: **p=0.001; -ConcA-D vs. +ConcA-D: **p=0.004; +ConcA+D vs. +ConcA-D: *p=0.025; all other comparisons not significant. **(G)** Quantification of average Ura7-mCherry event size (area, μm^2^). Statistics as in (E). -ConcA+D vs +ConcA+D: *p=0.04; -ConcA+D vs +ConcA-D: *p=0.013; all other comparisons not significant. **(H)** Quantification of filamentous Ura7-mCherry structures (aspect ratio (AR) ≥1.5) per cell. Statistics as in (E). -ConcA+D vs +ConcA-D: ***p=0.0003; -ConcA-D vs. +ConcA-D: ***p=0.0003; +ConcA+D vs. +ConcA-D: **p=0.002; all other comparisons not significant.

The synergistic effect of combining V-ATPase inhibition with glucose-induced disassembly fits a model in which Ura7 assembly responds to defects in vacuolar acidification and pH homeostasis. The V-ATPase helps regulate cytosolic pH by pumping protons into the vacuole (Martinez-Munoz and Kane, 2008), and yeast CTP synthase filamentation is triggered by drops in pH (Hansen et al., 2021). The “double hit” of inhibiting the proton pump with ConcA while forcing structural disassembly through glucose withdrawal would cause severe pH dysregulation. Glucose deprivation alone causes V-ATPase disassembly and rapid cytosolic acidification (Kane, 1995; Dechant et al., 2010); pre-treating with ConcA would prevent compensatory proton pumping during this transition. However, we have not directly measured pH under these conditions, and alternative mechanisms such as general metabolic stress signaling or ATP depletion could contribute. Future studies will be necessary to establish the causal link between V-ATPase dysfunction and CTP synthase assembly.

Collectively, our data establish spatial associations among the AP-3 vesicle coat complex, the V-ATPase, and CTP synthase, revealing a previously unrecognized organizational principle that links vesicular trafficking, pH regulation, and metabolic enzyme compartmentalization. While mechanistic details require further investigation, including direct pH measurements and biochemical study of protein interactions, these findings provide a conceptual framework for understanding this spatial coordination of metabolism and organellar function. Given that cytoophidium formation is conserved across eukaryotes (Liu, 2016; Noree et al., 2019), the connections identified here may reflect broader principles of how membrane trafficking influences metabolic enzyme assembly. Future investigations examining whether similar connections extend to other metabolic enzymes and occur in other organisms will determine the generality of these regulatory mechanisms.

## MATERIALS AND METHODS

### Yeast Strains and Culture Conditions

All *S. cerevisiae* strains used in this study (Table S2) were derived from the SEY6210 background (Robinson et al., 1988) and were constructed using standard techniques for yeast growth and genetic manipulation. Yeast strains were generated by one-step PCR-based integration (Longtine et al., 1998). Unless otherwise specified, cells were grown in rich medium (YPD: 1% yeast extract, 2% peptone, 2% glucose) at 26°C with shaking at 200 rpm.

### Bimolecular Fluorescence Complementation (BiFC) Analysis

BiFC reconstitutes a fluorescent Venus protein when complementary N-terminal (VN) and C-terminal (VC) fragments are brought into proximity by interacting or spatially adjacent proteins (Kerppola, 2008). To generate strains for BiFC analysis, we integrated genes encoding Venus C-terminal (VC) or Venus N-terminal (VN) fragments at the chromosomal loci of genes of interest. For BiFC experiments, MATα haploid strains expressing Apl6-VC or Vma10-VC were mated with MATa haploid strains expressing VN fusions to candidate proteins from the Bioneer VN collection (Sung et al., 2013) or custom-constructed VN fusions. Resulting diploids were sporulated, and haploid progeny expressing both VC and VN fusion proteins were selected and confirmed by genetic markers. For experiments with heterozygous diploids (Figures 1 and 2, and Supplemental Figure S2), MATα and MATa haploid strains carrying individual VC or VN fusions were mated, and the resulting diploid cells (expressing both tagged and untagged versions of each protein) were used directly for imaging. Prior to imaging, cells were grown to log phase (OD□□□ = 0.4–0.6), stained with CMAC-blue vacuolar marker (7-amino-4-chloromethylcoumarin; 50 μM final concentration) for 20 minutes at 26°C, washed, and immediately imaged. Note that CMAC-blue accumulates in the vacuolar lumen in nutrient-replete cells but is excluded from vacuoles and stains the cytosol in starved cells, as vacuolar acidification is reduced under nutrient deprivation.

### Nutrient Starvation Treatments

For starvation buffer experiments (Figures 3 and 4), cells were grown to log phase in YPD, collected by centrifugation at 3,000 × *g* for 3 minutes, washed once with sterile water, and resuspended at the original cell density in citrate-phosphate buffer (0.15 M, pH 6.0). Cells were then incubated with shaking at 26°C for 16–20 hours before imaging. For acute glucose withdrawal experiments (Figure 5), cells were grown to log phase in YPD, collected by centrifugation at 3,000 × *g* for 3 minutes, washed once with sterile water, and resuspended in YP medium (lacking glucose) at the original cell density. Cells were incubated with shaking at room temperature for 20 minutes, stained with CMAC-blue, and imaged.

### Concanamycin A (ConcA) Treatment

For ConcA treatment alone (Figure 5), cells were grown to log phase in YPD and concanamycin A (LC Laboratories) was added directly to the culture at a final concentration of 0.5 μM. Cells were incubated with shaking at room temperature for 90 minutes, stained with CMAC-blue, and imaged. For combined ConcA and glucose withdrawal experiments (Figure 5), cells were first treated with 0.5 μM ConcA for 90 minutes at room temperature as described above. After 90 minutes, cells were collected by centrifugation, washed once with water, and resuspended in YP medium (lacking glucose) supplemented with 0.5 μM ConcA. Cells were incubated for an additional 20 minutes at room temperature, stained with CMAC-blue, and imaged.

### Fluorescence Microscopy

All live-cell imaging was performed on a Nikon Ti2E inverted microscope equipped with a CSU-X1 Yokogawa spinning disk confocal scanner unit, OBIS 405, 488, 561, and 640 nm lasers, and an ANDOR iXon Ultra 512 × 512 EMCCD camera. Images were acquired using a Nikon Plan Apo λ 100× oil immersion objective (1.45 N.A.) and controlled with Micro-Manager 2.0 software. For z-stack acquisitions, three optical sections separated by 0.5 μm were collected and collapsed into 2D representations using maximum intensity projections in Fiji (ImageJ, NIH). Images were processed and assembled into figures using Fiji and Adobe Illustrator. All data are presented as mean ± SEM for biological replicate means. Statistical significance was assessed at α = 0.05 using GraphPad Prism software.

### Image Quantification and Statistical Analysis

All image quantification was performed using Fiji/ImageJ. For each experimental condition, ≥100 cells were analyzed per biological replicate, and all experiments were performed in biological triplicate. All datasets were first assessed for variance heterogeneity across conditions. Brown-Forsythe tests and F tests confirmed homogeneity of variance across conditions. Statistical comparisons were performed using one-way ANOVA with Tukey’s post-hoc test for experiments involving three or more groups, and unpaired t-tests for two-group comparisons. All analyses were performed on biological replicate means unless otherwise noted. Exact statistical tests and post hoc comparisons used for each dataset are specified in the figure legends and stated below.

#### BiFC quantification (Figures 1B,2B, 2E, S2B) and BiFC fluorescence quantification (Figures 1C, 2C, 2F, S2C)

Individual cells were manually segmented using region-of-interest (ROI) outlines, excluding cells with abnormally high diffuse autofluorescence indicative of compromised membrane integrity. Using a custom macro, cells were scored for the presence of ≥1 BiFC puncta per cell, and total fluorescence intensity values of puncta in each cell were recorded. Total puncta fluorescence per cell was divided by the median total puncta fluorescence measured in the His2 negative control from the same imaging session. Statistical comparisons of the average intensities from each replicate were performed using one-way ANOVA with Tukey’s post-hoc test.

#### Ura7 cytoophidia-vacuole colocalization (Figure 3D)

Multi-channel z-stacks were collapsed by maximum intensity projection and channels were separated for independent processing. Binary masks were generated for each channel using automated thresholding. To identify overlapping structures, the Image Calculator AND operation was applied to generate an intersection mask representing colocalized pixels. Objects in each mask were quantified using the Analyze Particles tool with size exclusion filters (0.05 μm^2^ to infinity) and circularity constraints to exclude noise. The percentage of Ura7-GFP cytoophidia colocalizing with Vph1-mCherry was calculated as: (number of objects in intersection mask / total number of Ura7-GFP objects) × 100. Statistical analysis was performed using unpaired t-test.

#### Ura7 filament-to-puncta ratio (Figure 4G)

Multi-channel images were separated and the Ura7-mCherry channel was processed using Otsu thresholding. Automated particle analysis was performed on binary masks with a size filter of 0.05 μm^2^ minimum to eliminate background noise. Ura7-mCherry structures were classified by aspect ratio (AR): AR ≥1.5 was categorized as filaments, AR <1.5 as puncta. Automated counts were manually verified against raw data. Statistical comparisons were performed using one-way ANOVA with Tukey’s post-hoc test.

#### Ura7 events per cell quantification (Figure 4H)

Total Ura7-mCherry events per cell were quantified in starved cells using the same thresholding and particle analysis approach described above for filament-to-puncta analysis. Events were defined as discrete structures with area ≥0.05 μm^2^ detected above threshold. The total number of events per cell was counted for each strain. Statistical comparisons were performed using one-way ANOVA with Tukey’s post-hoc test.

#### Ura7 assembly quantification following V-ATPase inhibition (Figures 5E-H)

For quantification of Ura7 structures (puncta and filaments), the Ura7-mCherry channel was thresholded using the Otsu method. The threshold value determined for wild-type mock-treated cells was applied uniformly to all experimental conditions to ensure consistent detection. Particle analysis was performed to quantify the number, area, integrated density, and aspect ratio of Ura7 structures. Filamentous structures were defined as objects with AR ≥1.5 and area ≥0.5 μm^2^. Statistical analysis was performed using one-way ANOVA with Tukey’s post-hoc test.

#### Classification of Ura7 structures

Following established criteria (Noree et al., 2010; Barry et al., 2014), Ura7 structures were classified based on morphology. Structures <0.75 μm in length are defined as puncta (foci), representing nucleation sites or smaller assemblies, while structures ≥0.75 μm are defined as filaments (cytoophidia). For automated quantification of filament-to-puncta ratios, we used aspect ratio (AR) as a proxy for elongation, with AR ≥1.5 corresponding to filamentous structures and AR <1.5 corresponding to punctate structures. This approach was validated by manual inspection confirming that structures with AR ≥1.5 consistently represented elongated filaments rather than circular puncta.

## ACKNOWLEDGEMENTS

We thank Michael McMurray (University of Colorado Anschutz School of Medicine) for supplying VN-fusion yeast strains.

## SUPPLEMENTAL FIGURE LEGENDS

**Figure S1. BiFC screening strategy and candidate validation**

**(A)** Schematic diagram of the AP-3 complex. **(B)** BiFC screening workflow schematic. MATα haploid cells expressing Apl6-VC were mated with MATa cells expressing VN fusions. Diploid cells were examined by confocal microscopy. This approach systematically assessed the 98 candidates from Table S1 for spatial proximity to AP-3. Representative confocal images of cells expressing Apl6-VC paired with VN fusions to His2 (negative control) versus Apl5 (positive control) **(C)** and to the candidate proteins in Table S1 that produced BiFC puncta **(D)**. Scale bar, 5 μm.

**Figure S2. The CTP synthase homolog Ura8 associates with AP-3 by BiFC**

(A) Representative confocal images of cells expressing the indicated VN/VC fusions. Scale bar, 5 μm. Quantification of **(B)** BiFC-positive cells **p=0.004; ns, not significant (Ura8-VN His2-VC vs. Ura8-VN Apl6-VC, p=0.077; Ura8-VN His2-VC vs. Ura8-VN Apl6-VC, p=0.086) and **(C)** total BiFC fluorescence. ns, not significant (p=0.229). Graphs and statistics as in Figure 1B,C.

