## Supplementary material for "AP-3 and the V-ATPase Modulate CTP Synthase Assembly Through Spatial Association at the Yeast Vacuole": Figure S1

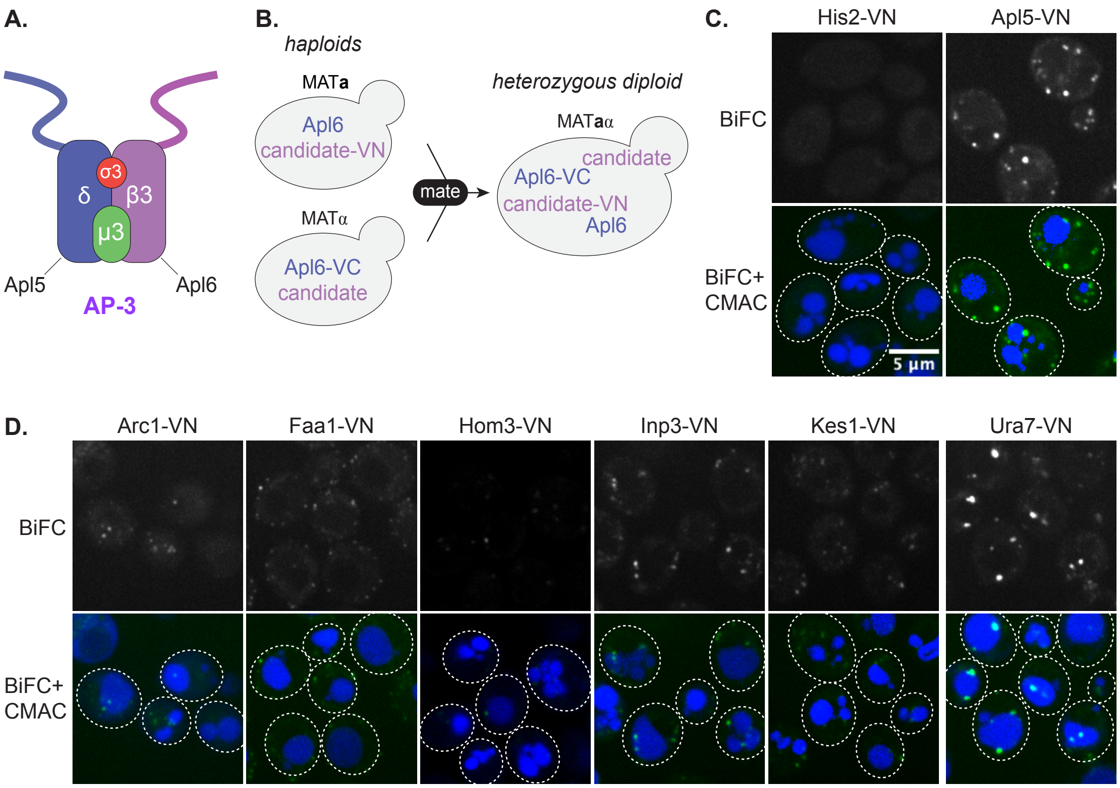


**Supplemental Figure S1. BiFC screening strategy and candidate validation**

**(A)** Schematic diagram of the AP-3 complex. **(B)** BiFC screening workflow schematic. MATα haploid cells expressing Apl6-VC were mated with MATa cells expressing VN fusions. Diploid cells were examined by confocal microscopy. This approach systematically assessed candidates the 98 candidates from Table S1 for spatial proximity to AP-3. Representative confocal images of cells expressing Apl6-VC paired with VN fusions to His2 (negative control) versus Apl5 (positive control) **(C)** and to the candidate proteins in Table S1 that produced BiFC puncta **(D)**. Scale bar, 5 μm.
