## Supplementary material for "AP-3 and the V-ATPase Modulate CTP Synthase Assembly Through Spatial Association at the Yeast Vacuole": Figure S2

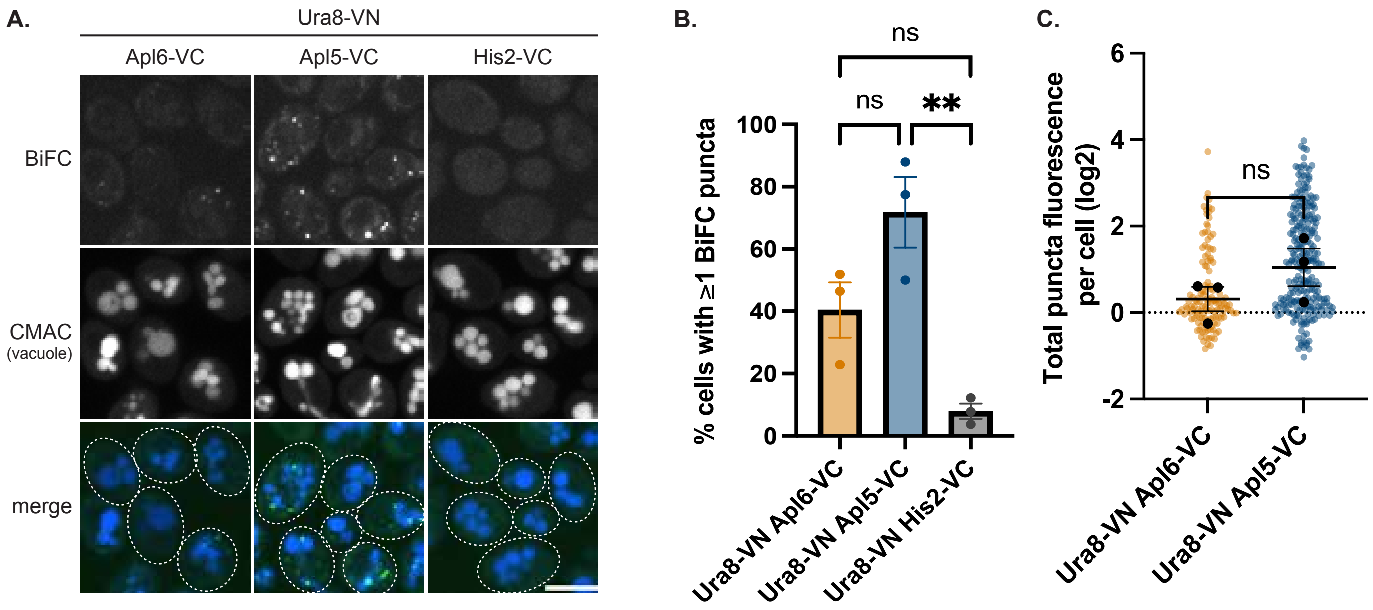


**Figure S2. The CTP synthase homolog Ura8 associates with AP-3 by BiFC**

**(A)** Representative confocal images of cells expressing the indicated VN/VC fusions. Scale bar, 5 µm. Quantification of **(B)** BiFC-positive cells **p=0.004; ns, not significant (Ura8-VN His2-VC vs. Ura8-VN Apl6-VC, p=0.077; Ura8-VN His2-VC vs. Ura8-VN Apl6-VC, p=0.086) and **(C)** total BiFC fluorescence. ns, not significant (p=0.229). Graphs and statistics as in Figure 1B,C.
